# CG methylation covaries with differential gene expression between leaf and floral bud tissues of *Brachypodium distachyon*

**DOI:** 10.1101/024935

**Authors:** Kyria Roessler, Shohei Takuno, Brandon S. Gaut

## Abstract

DNA methylation has the potential to influence plant growth and development through its influence on gene expression. To date, however, the evidence from plant systems is mixed as to whether patterns of DNA methylation vary significantly among tissues and, if so, whether these differences affect tissue-specific gene expression. To address these questions, we analyzed both bisulfite sequence (BSseq) and transcriptomic sequence data from three biological replicates of two tissues (leaf and floral bud) from the model grass species *Brachypodium distachyon.* Our first goal was to determine whether tissues were more differentiated in DNA methylation than explained by variation among biological replicates. Tissues were more differentiated than biological replicates, but the analysis of replicated data revealed high (>50%) false positive rates for the inference of differentially methylated sites (DMSs) and differentially methylated regions (DMRs). Comparing methylation to gene expression, we found that differential CG methylation consistently covaried negatively with gene expression, regardless as to whether methylation was within genes, within their promoters or even within their closest transposable element. The relationship between gene expression and either CHG or CHH methylation was less consistent. In total, CG methylation in promoters explained 9% of the variation in tissue-specific expression across genes, suggesting that CG methylation is a minor but appreciable factor in tissue differentiation.

## Introduction

The term ‘epigenetics’ refers to processes beyond (*epi*-) genetics and, more concretely, to heritable chromosomal modifications that have the potential to vary during development and stress [1,2]. Epigenetic modifications include histone variants but also DNA methylation. In plants, the methylation of cytosines occurs in three sequence contexts: CG, CHG and CHH, where H = A, C or T. All three contexts are usually methylated in repetitive sequences, which serves to limit the transcription and proliferation of transposable elements (TEs) [3]. Genes are often also methylated, too, but typically only in the CG context [4-6]. The function of this gene-body methylation (gbM) is not yet clear, but potential functions include the exclusion of histone H2A.Z [7], control of aberrant intragenic expression [8], protection from transposable element (TE) insertion [9], and facilitation of intron-exon splicing [10-12].

DNA methylation has long been hypothesized to have a direct effect on gene regulation during development [13,14]. With the growing availability of single base resolution methylation data, like bisulfite sequencing (BSseq) data, this hypothesis has been tested directly. In humans, for example, DNA methylation varies dramatically throughout development, and this variation is often correlated with gene expression [15-18]. In plants, the available evidence suggests that methylation levels vary for highly specialized tissues, such as the endosperm and the pollen vegetative nucleus [19-22], and that this methylation variation likely contributes to genetic imprinting and trans-generational silencing of TEs [23-25].

Outside of these few specialized tissues, a clear picture has not yet emerged as to whether methylation commonly varies among plant tissues and, if so, whether methylation variation contributes to tissue-specific gene expression (GE). Some evidence suggests that most plant tissues do not vary substantially in DNA methylation. For example, genome-wide profiling in rice (*Oryza sativa* L.) identified few DNA methylation differences between shoot and root, and only a few additional differences in CHH methylation between these two tissues and the embryo [22]. Moreover, a survey of several *A. thaliana* accessions found that tissue-specific variation in methylation was much less pronounced than genetic variation, leading the authors to conclude that “…DNA methylation is less dynamic than gene expression patterns in plants and only plays a role during specific stages of development or cell type, such as companion cells” [26].

In contrast to these studies, there is some emerging evidence that differential methylation may play a role in tissue specific GE. For example, researchers have detected ∼2000 differentially methylated regions (DMRs) among four soybean tissues, and a subset of these DMRs correlate with tissue-specific GE of ∼60 genes [27]. Similarly, analysis of tissue-specific DNA methylation patterns in *Sorghum bicolor [28]*, *Populus tichocarpa* [29] and maize (*Zea mays* ssp. *mays*) [30] hint that epigenetic variation among vegetative tissues correlates with tissue-specific expression. However, not all of these studies have measured methylation at single-base resolution, which greatly limits the ability to draw firm conclusions; the number of contrasts of methylation between plant tissues is growing, but such studies remain rare.

Methodological differences among studies have also made conclusions difficult. For example, few methylation studies have employed biological replication, and thus it is usually unclear whether methylation variation between tissues exceeds the statistical variation expected from within a single sampled tissue. Even when the data are at single-base resolution, studies have used different summaries of the data as the basis for inferences, and this has caused confusion. Some studies have focused on summarizing methylation for genomic features like genes and TEs [31,32]. Other studies have focused on DMRs as a summary of the data. DMRs were initially defined as stretches of DNA sequence for which methylation differences between samples were higher than expected at random [16]. While the initial definition of DMRs is straightforward and meaningful, more recent studies have used empirical means, such as sliding windows, to identify DMRs, and these empirical definitions vary from study to study [27,33]. As a result, the interpretation and meaning of DMRs varies among studies, compromising the value of inferences.

This study is focused ultimately on the question of whether DNA methylation and GE covary between tissues. To that end, we have measured both DNA methylation and gene expression between two tissues (leaves and floral buds) of *Brachypodium distachyon* (brachypodium), a grass species that has served as a model for genomic studies [34]. While our ultimate goal is to assess methylation and GE, our proximal goals include an empirical assessment of the effects of replication both on inferring methylation differentiation between tissues and on the impact of summary methods (i.e., DMRs vs. single-base metrics) on inferences. Overall, we find the two tissue samples to be significantly different in DNA methylation patterns, but we also find that the false positive rate without replication is high (>50%). In all respects, DMRs are less useful than single-base or regional measures in our empirical analyses. Altogether, we find that CG methylation and GE covary between tissues, explaining up to 9% of variation in gene expression.

## Results and Discussion

### DNA Methylation within and between tissue samples

To assess methylation variation, we utilized BSseq data from a previous study [35] that generated reads from three biological replicates of two tissues: leaf and immature flower buds. We denoted the leaf replicates as L1, L2 and L3 and the floral bud replicates as F1, F2 and F3. The data had conversion error rates of <1.3% for each replicate [35](S1 Table). Following the previous study, we mapped BSseq reads to the *B. distachyon* genome and tallied only uniquely mapping reads. Each replicate yielded ∼15X of mapped coverage, such that each tissue had ∼45X coverage per base, on average [35].

To our knowledge, no plant DNA methylation papers have assessed whether tissue-specific variation exceeds that expected from proper biological replication. To assess this question, we first tested for a signal of differentiation between two BSseq datasets at single nucleotide sites, which we call <u>D</u>ifferentially <u>M</u>ethylated <u>S</u>ites (DMSs). To identify DMSs, we required a minimum coverage of 3 reads for each site in each tissue and then applied Fisher’s Exact Test (FET) [16] (see Methods). There were many DMSs between two biological replicates from the same tissue. For example, there were 218,631 DMSs between L1 and L2 and an average of ∼250,000 DMSs between two leaf replicates (Fig. 1A). However, DMSs were more abundant between replicates from different tissues, with an average of ∼324,000 differential sites. (Fig. 1A). The average number of DMSs was significantly higher for between-tissue vs. intra-tissue comparisons (permutation, *p*<0.01), indicating that the tissue samples were significantly differentiated.

**Fig. 1:**
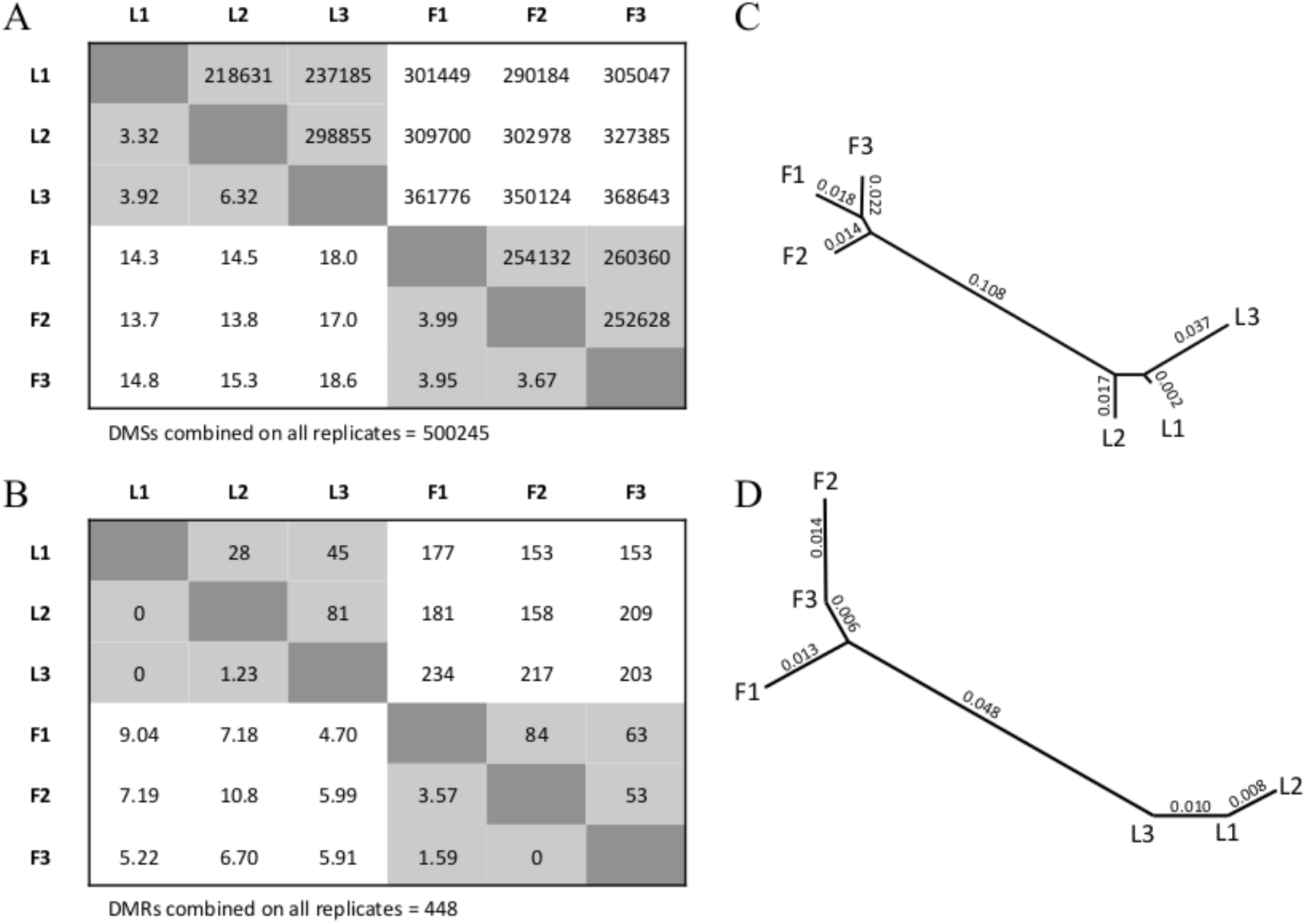
The inferred number of DMSs and DMRs between replicates. A) The upper matrix reports the number of DMSs between two BSseq replicates. The lower matrix percentage of DMSs that map to the same location as the 500,245 DMSs inferred from the combined data set. B) A neighbor-joining phylogeny representing the relationship among the six BSseq samples, based on distances defined by the lower matrix in A. C) The upper matrix reports the number of DMRs between two BSseq replicates. The lower matrix provides the percentage of DMRs that overlap with the 448 DMRs inferred from the combined data set. D) A neighbor-joining phylogeny representing the relationship among the six BSseq samples, based on distances defined by the lower matrix in C.

We also inferred DMSs by combining the three replicates within each tissue and then comparing the combined leaf data (L1+L2+L3) to the combined floral data (F1+F2+F3). Using the combined replicates from each tissue, FET analyses identified 500,245 DMSs. Following a previous study [36], we assumed these 500,245 DMSs to be our best estimate of “true” DMSs between tissues and found that this true set rarely overlapped in location with DMSs that were identified between replicates within a tissue; typically <5% of within-tissue DMSs overlapped with the true set (Fig. 1A). In contrast, the overlap was more significant, at 15.6% on average, between the true set and DMSs identified between replicates from different tissues (Fig. 1A). These percentages define genetic distances between two BSseq replicates that can then be used for clustering analyses. A neighbor-joining analysis clearly separated replicates from different tissues (Fig. 1B), further supporting the contention that the two tissue samples differ in DNA methylation beyond that expected from sampling.

It is interesting to note, however, that comparisons between single-replicates yield high false-positive rates (FPRs). For example, the comparison of L3 and F3 replicates yielded 368,643 DMSs (Fig. 1A). Of these, 92,989 (or 18.6%) overlapped with the set of true DMSs; hence, 81.4% of the DMSs identified between these two replicates were not supported by the larger, combined data set. In other words, had we relied on a single replicate from each tissue for this study, >80% of our inferences would have been incorrect relative to more extensive data. In theory, the high FPR can be reduced either by increasing the stringency of the FET or by adjusting the FET for multiple-tests, but these adjustments do not help in this case. For example, when we focused on the L3 vs. F3 comparison and applied a false discovery rate (FDR) correction at *q* < 0.05, we found 10,435 DMSs compared to 368,643 without FDR adjustment (Fig. 1A). However, none of these DMSs overlapped with the true set, yielding an FPR of 100%.

A potential advantage of studying DMRs, as opposed to DMSs, is that they summarize signals over contiguous sites, and it is thus possible that they reduce the FPR. As we have noted, the definition of a DMR varies widely among studies; here we focused on the original definition to define DMRs as a region of non-random differentiation between samples [16]. To determine expectations under ‘randomness’, we permutated cytosine methylation states throughout the genome (see Materials & Methods); permutations indicated that ≥ 5 DMSs in a row were 1.3% of those expected at random (S1 Fig.). Accordingly, we defined a DMR as ≥ 5 DMSs in a row that had a consistent direction of methylation bias (i.e., hypomethylated in one or the other tissue).

With this definition, we again observed more DMRs between tissues (187, on average) than within tissues (59, on average)(Fig. 1C), and this difference was again statistically significant (permutation, *p*<0.01). By combining data (L1+L2+L3 vs. F1+F2+F3), we inferred a ‘true’ set of 448 DMRs with an average size of 38.4 bp, a minimum length of 5.00 bp and a maximum length of 522 bp. On average, 1.0% of within-tissue DMRs overlapped with the true set, whereas 6.9% of between-tissue DMRs overlapped with the true set on average. These distances again separated the tissues in clustering analyses (Fig. 1D), verifying significant tissue differentiation. However, the between-tissue FPR based on single BSseq replicates was consistently higher for DMRs than for DMSs, with a *minimum* FPR of 89.2%. In other words, ∼80% or more of our DMR inferences were incorrect for contrasts between single replicates relative to the more extensive dataset.

This raises the question as to why the FPR is so high and whether our observations are unique. To answer the latter question, to our knowledge only one other study of plants has used biological replication for BSseq data to compare methylation between *Arabidopsis thaliana* and *A. lyrata* [37]. (At least one other paper replicated their control but not experimental samples [38]) In the *Arabidopsi*s paper, the authors used replication to help filter the number sites for testing, thus reducing the multiple test problem and increasing statistical power. They did not, however, explicitly report on the level of within vs. between tissue differences. To address the former question, the FPR may be high for technical, statistical and/or biological reasons. Technically, BSseq data are subject to conversion error, but conversion errors are unlikely to explain our observations because coverage is high and conversion error is low. Statistically, it is easy to envision that the FET may signal numerous false-positives, but the FPR remains high for DMSs when the FET is FDR corrected [39], as noted above. Finally, biological variation among replicates may contribute to the FPR, both because tissue samples are likely to include mixtures of different cell types that vary in proportion among replicates [40] and also because it is likely that there is heterogeneity in methylation levels even among cells of a single type [41]. We thus suspect, but cannot prove, that the largest contribution to variation among replicates is biological in origin.

Our FPR calculations deserve two further comments. First, the FPR calculations rely on the assumption that the set of ‘true’ DMSs and DMRs are defined by our analysis of combined data. This assumption cannot hold fully because there must be false-positives in the combined data, but the FPR rate of the combined data are difficult to assess. Second, we note that as levels of methylation differentiation become more pronounced, the signal:noise ratio will also increase. Thus, our data reflect the importance of replication for contrasts between tissues in the same species; however, it may not be as useful to replicate data that are designed to summarize broad-scale differences in methylation patterns between distantly related species [31,32].

### Differences in methylation patterns between tissues

Given that we found tissue-specific differentiation between leaves and floral buds, we sought to categorize the pattern of methylation differences, both in terms of cytosine contexts and genomic locations. For these results, we based all analyses on combined leaf (L1+L2+L3) and floral (F1+F2+F3) data, following [36].

Our first finding was that the two tissue samples were more similar than different in their methylation patterns. To reach this conclusion, we identified sites with conserved methylation between the two tissues – i.e., Conserved Methylated Sites (CMSs). To be a CMS, a site required the support of a binomial test [5] at a *p*-value of 0.05 for a site in both samples. Overall, we found that 18,780,682 cytosines were methylated in both tissues (Fig. 2A; Table 1), representing 18.7% of the 100,229,480 genomic cytosines in the proper context for methylation (i.e., CG, CHG or CHH). Among CMSs, most (62.7%) were in the CG context, with an appreciable minority in the CHG context (29.8%) and relatively few in the CHH context (7.5%). Overall, the set of CMSs and DMSs were mutually exclusive, and there were 37-fold more CMSs (Fig. 2A; Table 1). Other studies have also found more similarities than differences among plant tissue samples [22,27].

**Fig. 2:**
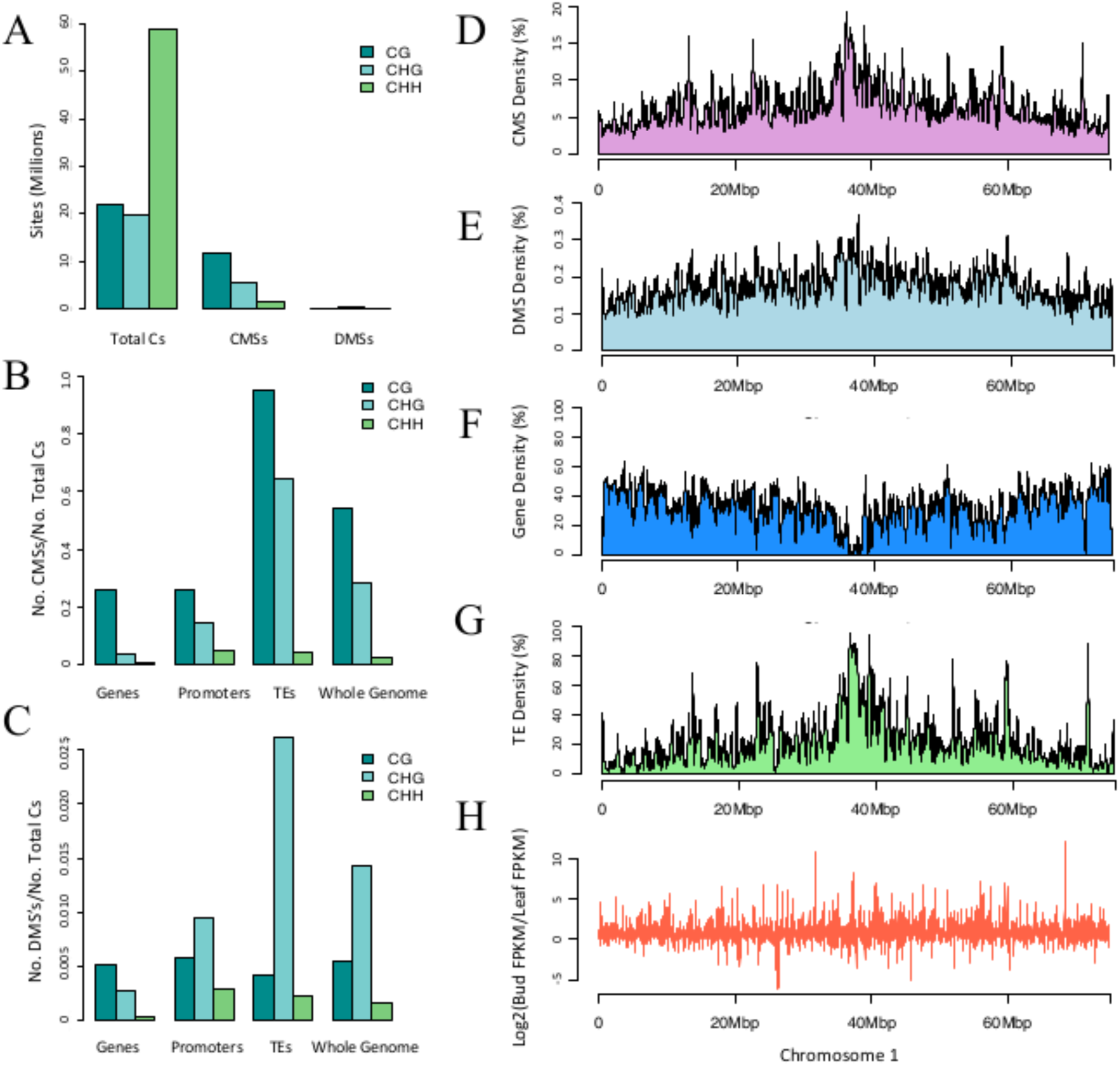
Context, direction, and regions of CMSs and DMSs. A) The number of sites in the correct context for methylation throughout the genome (Total Cs), along with the number of CMSs and DMSs in context. B) The proportion of CMSs relative to cytosines in the correct context for Genes, Promoters, TEs and the Whole Genome. C) The proportion of DMSs relative to cytosines in the correct context for Genes, Promoters, TEs and the Whole Genome. Note the difference in the scale of the *y*-axis between panels B and C. D to F) The graphs show the CMS, DMS, gene and TE density along chromosome 1. Density was measured within a 50kb sliding window for smoothing. H) Differential gene expression plotted along the physical length of chromosome 1. The other chromosomes are represented in S2 Fig.

**Table 1:** The number of potential methylation sites, DMSs and CMSs in each of three sequence contexts (CG, CHG and CHH) throughout the entire brachypodium genome and also for three features separately (Genes, Promoters and TEs). Numbers in parentheses represent the percentage of sites in context that are methylated.

| Region | CG | CHG | CHH | Total |
| --- | --- | --- | --- | --- |
| Potential methylation sites |  |  |  |  |
| Genic | 5,945,276 | 6,261,258 | 17,109,118 | 29,315,652 |
| TE | 7,066,299 | 5,524,931 | 17,168,841 | 29,760,071 |
| Promoter | 2,201,304 | 1,833,675 | 5,378,012 | 9,412,991 |
| Whole Genome | 21,814,767 | 19,722,162 | 58,692,551 | 100,229,480 |
| Differentially Methylated Sites (DMSs) |  |  |  |  |
| Genic | 30,396<br>(0.51) | 17,059<br>(0.27) | 6,009<br>(0.04) | 53,464<br>(0.18) |
| TE | 29,981<br>(0.42) | 144,779<br>(2.62) | 39,990<br>(0.23) | 214,750<br>(0.72) |
| Promoter | 12,658<br>(0.58) | 17,276<br>(0.94) | 15,500<br>(0.29) | 45,434<br>(0.48) |
| Whole Genome | 120,744<br>(0.55) | 282,440<br>(1.43) | 9,7061<br>(0.17) | 500,245<br>(0.50) |
| Conserved Methylated Sites (CMSs) |  |  |  |  |
| Genic | 1,537,550<br>(25.86) | 216,844<br>(3.46) | 88,467<br>(0.52) | 1,842,861<br>(6.29) |
| TE | 6,736,090<br>(95.33) | 3,560,407<br>(64.44) | 663,269<br>(3.86) | 10,959,766<br>(36.83) |
| Promoter | 568,236<br>(25.81) | 265,069<br>(14.46) | 232,488<br>(4.32) | 1,065,793<br>(11.32) |
| Whole Genome | 11,776,244<br>(52.98) | 5,590,834<br>(28.35) | 1,413,604<br>(2.41) | 18,780,682<br>(18.74) |

Our second finding was that most variation between tissues occurred at CHG sites. Cytosines were most commonly methylated in the CG context, but 56.5% (282,440 sites) of DMS sites occurred in the CHG context (Table 1; Fig. 2A). To investigate further, we estimated the DMS ‘rate’ by comparing the observed number of DMSs to the available number of cytosines in a particular context. For example, there were 19,722,162 cytosines in the CHG context throughout the genome and a total of 282,440 DMSs, yielding a rate of 1.43% (Table 1). In contrast, CG and CHH methylation had lower rates, at 0.55% and 0.17%, respectively (Table 1). Interestingly, the direction of DMSs was biased, because 57% were methylated in floral buds but not leaf, representing a deviation from the expectation of equality (binomial, p<10^−15^).

Our observation that variability between tissues was highest at CHG sites is similar to comparisons among rice tissues [22], which found that the highest methylation variation was between embryo and endosperm in the CHG context. Similarly, in *Arabidopsis* species tissue-specific differences were attributable to CHH and CHG methylation changes within DMRs [37]. However, CG methylation varies more than CHH or CHG methylation among tomato developmental stages [42] and also between generations of *A. thalian*a mutation accumulation lines [39]. Thus, the principle context of DNA methylation variability varies either as a function of species or perhaps the tissues sampled.

Having assessed the effect of context, we shifted our attention to three genomic features of interest: genes, promoters and transposable elements (TEs). Among the three features, the set of 68,264 non-genic, annotated TEs had the highest CMS rates (Fig. 2B), as was expected from previous studies of plant genomes [39,43], with methylation level of 95.3% at CG sites and 64.4% at CHG sites (Table 1; Fig. 2B). That said, TEs also had the highest DMSs rates, at 2.62% in the CHG context (Table 1; Fig. 2C). CHH methylation levels were low (<5%) throughout TEs, as noted previously for the entire brachypodium genome [35]. In contrast to TEs, the 26,072 annotated genes had the lowest DMS rate at 0.18% (Table 1; Fig. 2C), but this low rate may reflect the fact that genes were primarily methylated in the CG context, which had the lowest DMS rates. Promoter regions, which were defined as 1.0 kb 5’ upstream of the 26,072 genes, had noticeably higher levels of conserved CHG methylation between tissues (at 14.5%) than genes (3.46%), but were similar to genes in most other respects (Fig. 2; Table 1).

Given that CMSs and DMSs were especially prominent in TEs, it was not surprising that the distribution of CMSs and DMSs across chromosomes mimicked the density of TEs (Fig. 2 and S2 Fig.), but there was no obvious correlation between CMSs and DMSs with gene density (Fig. 2G and S2 Fig). Altogether, the analysis of single sites paints a clear picture: most methylation occurred in TEs and most variation between tissues was within TEs in the CHG context.

Finally, we examined the pattern and location of the 448 DMRs identified between tissues to assess whether they paralleled results based on single sites. First, 65% of DMRs were hypo-methylated in floral buds (p < 0.01), verifying increases in overall methylation in this tissue. Second, although most DMSs were found in the CHG context (Fig. 2C), we found that 67% of the DMSs within all of our DMRs were sites in the CHH context. This observation suggests that there may be a spatial (clustered) context to the mechanisms that underlie CHH differences between tissues, consistent with the observation in maize that CHH sites tend to be clustered [43]. Finally, the location of DMRs was biased: 39% of DMRs were found in unannotated regions of the genome, but 31% were found within TEs, 17% within genes and 13% within promoter regions. Given that the total number of cytosine sites within TEs and within genes was similar, at ∼29 million bases (Table 1) each, the lower percentage in genes again indicates that genic methylation is more highly conserved between tissues than methylation of TEs.

### Methylation & Gene Expression

The primary goal of this paper is to determine whether methylation differentiation between tissues covaries with GE. The idea that GE and methylation covary traces back to the origin of epigenetics [44] and seems to be upheld by weak signals from plant data [27,42].

#### Gene expression data

To measure GE, we generated RNAseq data from leaf and floral tissues, using the same three plants and samples (biological replicates) that were used to generate BSseq data (see Methods). Each of the replicates had > 12 million RNAseq reads that mapped uniquely to the *B. distachyon* genome (S2 Table). Out of 26,552 annotated protein-coding genes, we retained 26,072 that did not overlap with annotated TEs, of which 19,956 had evidence of expression in at least one tissue, as determined by a cutoff of FPKM > 0.02 (see Materials and Methods). Second, we identified differentially expressed genes between tissues at an FDR of *q* < 0.01 (S3 Fig.). A total of 7,704 genes were significantly differentially expressed between leaf and floral tissue; these exhibited no obvious clustering by chromosomal position (Fig. 2H and S2 Fig.). GO analyses of differentially expressed genes suggested enrichment for functions in membrane and microtubule development (S3 Table).

#### GE and DMRs

Since many studies have focused on DMRs (rather than annotation features) to assess correlations with GE, we began by testing for associations between GE and DMRs. If DMRs influence differential GE, we hypothesized that DMRs should be enriched around differentially expressed genes. To test this hypothesis, we measured the distance (in bp) between a DMR and the closest differentially expressed gene. We then tested whether the observed average distance from DMRs to genes was smaller than expected at random, as tested by permutation (see Materials & Methods). We found that on average a DMR was 18,710 bp from a differentially expressed gene, which was not significantly smaller than the random expectation of 17,572 bp (*p*=0.82; S4 Fig.). Based on this analysis, there is no evidence to suggest that DMRs are enriched near differentially expressed genes, as one might expect if DMRs help drive tissue-specific expression on a genome-wide scale.

Thinking that we may have missed an important signal by focusing on the entire genome, we delved into the three genomic features separately. For each feature, we focused on DMRs that were hyper-methylated in one vs. the other tissue. For example, we tallied DMRs within 25 genes that were hyper-methylated within floral buds. For this set of 25 genes, we predicted lower GE in floral than leaf tissue. Similarly, for the 19 genes that had a methylated DMR in leaf but not floral tissue, we predicted higher GE in leaf. These predictions were not upheld by the data, however (Fig. 3A). In fact, the average level of differential expression did not vary among genes that had a hyper-methylated DMR in floral bud, a hyper-methylated DMR leaf or no DMR whatsoever (Fig. 3A). We repeated this analysis for promoter regions of 1.0 kb 5’ upstream of genes, and again found no discernible pattern (Fig. 3B). Finally, because the methylation of TEs may effect the expression of nearby genes [3,45], we also examined DMRs within annotated TEs closest to a gene. Again, there was no signal (Fig. 3C). While the lack of signal may reflect low sample sizes, the presence of DMRs did not correlate with differential GE between the two tissues.

**Fig. 3:**
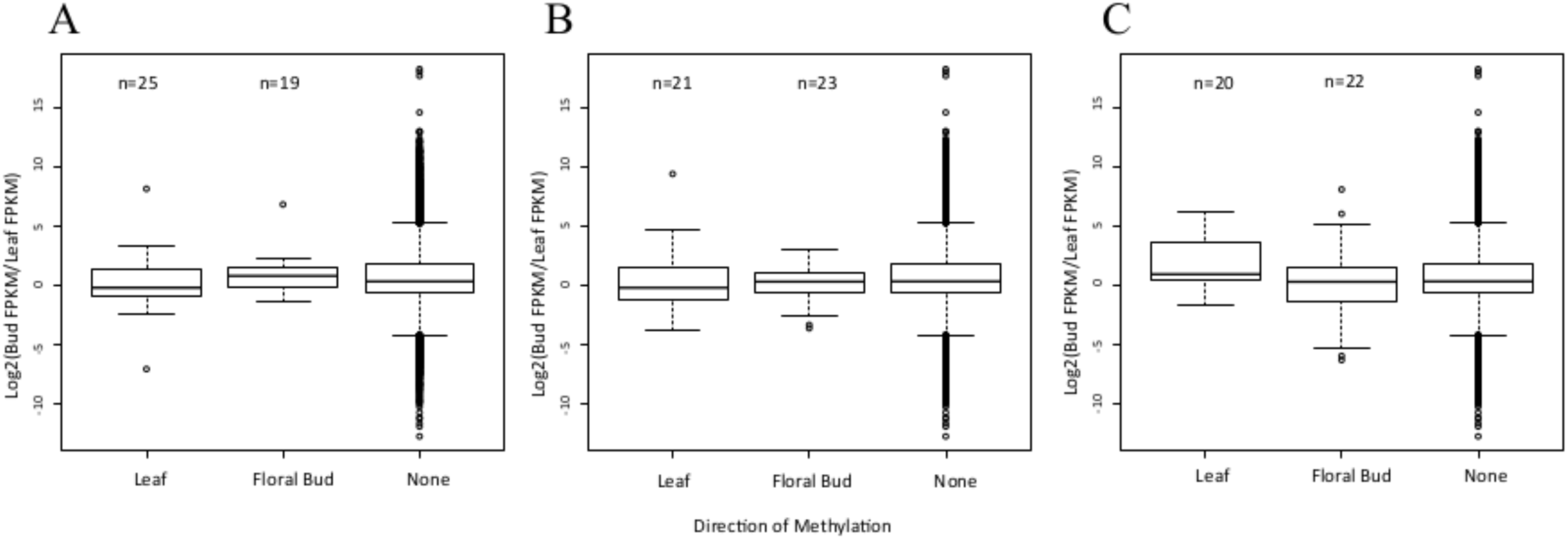
Gene expression with respect to DMRs and their direction. A) A graph of the distribution of gene expression when a DMR is located within a gene and hypermethylated in the Leaf or Floral Bud, or when there is no DMR in the gene (None). For the 25 genes hypermethylated in leaf, we predicted positive values on the *y*-axis, signaling higher expression in floral bud, but no bias was detected. For the 19 genes hypermethylated in floral bud, we predicted negative values on the *y*-axis, signaling higher expression in leaf, but again no bias was detected. B) The same graph of differential expression when the gene contains a DMR in its a promoter region. Again, there are no detectable biases in the direction of gene expression relative to genes that do not contain a DMR in their promoter region. C) A graph of differential gene expression when the TE nearest to a gene has a DMR that is hypermethylated in leaf, flower or no (None) DMR. For all graphs, the box plots represent the median, first, and third quartile. The whiskers represent the minimum and maximum, The numbers above the graph refer to sample size in each category.

#### The proportion of converted reads

To investigate covariation between GE and methylation more thoroughly, we turned to a measure of DNA methylation that summarizes the proportion of non-converted reads over the total number of reads at cytosine residues in the proper contexts (CG, CHG or CHH) [32]. This measure, which we call *prop*_*C*_, can be applied to the entire genome, to specific genomic features or to specific contexts (e.g., *prop*_*CG,*_ *prop*_*CHG,*_ *prop*_*CHH*_). For example, over the entire genome, *prop*_*c*_ was estimated to be 0.1815 for leaf tissue and 0.1823 for floral bud tissue throughout the entire genome, suggesting again (very) slightly higher levels of methylation in floral bud tissue. The *prop* measures provide an estimate of the methylation level for a region, but without a corresponding measure of significance. We focus on the use of these measures for the remainder of our analyses.

#### GE and Genic Methylation

To better understand patterns of methylation within genes, we first assessed the relationship among *prop*_*CG*_, *prop*_*CHG*_, and *prop*_*CHH*_ within a tissue, using correlation analyses. In brief, all are significantly correlated with one another, with *r* values ranging from 0.43 to 0.61 (Table 2). However, *prop*_*CG*_ and *prop*_*CHG*_ were positively correlated in a somewhat striking pattern: CHG methylation was often present but rarely higher than CG methylation (i.e., in only 3,424 of 26,072 genes) (Fig. 4A). This observation reaffirms that methylation in the CG context is predominant for genes [4,5,46] but also illustrates that genic methylation is not limited to the CG context [47].

**Fig. 4.**
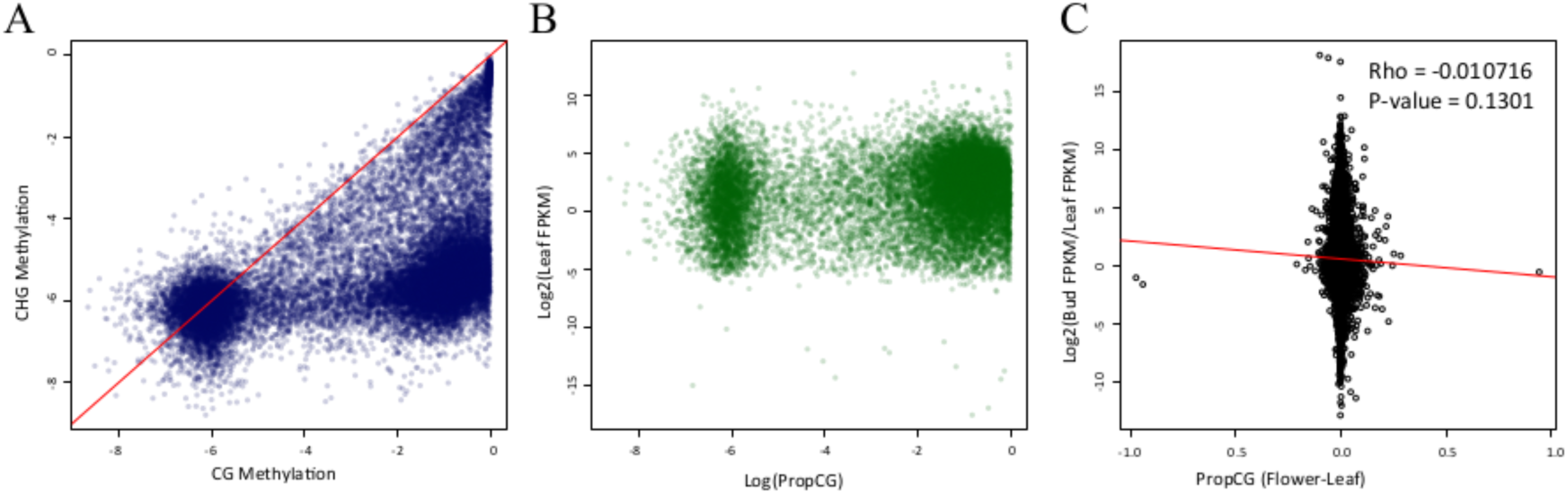
Methylation patterns within genes. A) The correlation between *prop*_CG_ and *prop*_CHG_ between genes for leaf tissue (*r* = 0.5977; *p* < 2.2e-16); floral bud tissue is not shown but the relationship is essentially identical. Methylation is plotted on a log scale. B) A comparison of propCG, on a log scale, and gene expression (FPKM) on a log2 scale within leaf (*r* = 0.2867; *p* <2.2e-16); again, floral bud tissue is not shown but essentially identical. C) A comparison of differential gene expression [log2fold (flower/leaf)] vs. the difference in *prop*_CG_ between leaf and floral bud tissue.

**Table 2:** Spearman correlation coefficients between *prop* values within a tissue. The p-values of all coefficients are < 2.2 e-16 and significant after sequential Bonferroni correction.

|  |  | Flower |  | Leaf |  |
| --- | --- | --- | --- | --- | --- |
|  |  | CHG | CHH | CHG | CHH |
| **Genes** | CG | 0.5976<sup>1</sup> | 0.4393 | 0.5977 | 0.4416 |
|  | CHG | --- | 0.6089 | --- | 0.6045 |
| **TEs** | CG | 0.3665 | -0.1513 | 0.3558 | -0.1743 |
|  | CHG | --- | 0.0639 | --- | 0.06036 |
| **Promoters** | CG | 0.7970 | 0.5567 | 0.7495 | 0.5436 |
|  | CHG | --- | 0.7173 | --- | 0.4293 |

Given that CG methylation is the primary component of genic methylation, we compared *prop*_*CG*_ to GE within a tissue. Previous work has shown that the relationship between GE and gene body methylation is complex [5]. In general, methylated genes have intermediate levels of expression, such that hypo-methylated genes are both more-highly and less-highly expressed than hyper-methylated genes [32,46,48]. As expected, GE and genic methylation were correlated within tissues (*r*=0.287; *p* < 2.2e-16; Table 2), but in a complex pattern (Fig. 3B). *prop*_*CHG*_ and *prop*_*CHH*_ were also correlated with GE but at lower levels (*r*=0.046, *p*=1.21×10^−13^ and *r*=0.073, *p*<2.2×10^−18^).

Lastly, we compared differential methylation to differential GE between tissues, focusing on either all of the 19,956 genes or just the 7,704 that were significantly differentially expressed. Differential GE and methylation were not correlated with *prop*_C_, *prop*_CHH_ or *prop*_CHG_ (Table 3; Fig. 3C) but were correlated between CG methylation and differential expression of the subset of 7,704 genes (Table 3). This significant correlation was negative, indicating that higher gene expression covaries with lower methylation levels. Note that the correlation, while significant, had a low absolute value (*r* = -0.0393; Table 3), suggesting that methylation differences explain at best a small proportion (3.9%) of the variance in GE between tissues. To sum: on a genome-wide scale, we uncovered moderate evidence that CG methylation and differential GE covary within genic regions.

**Table 3:**
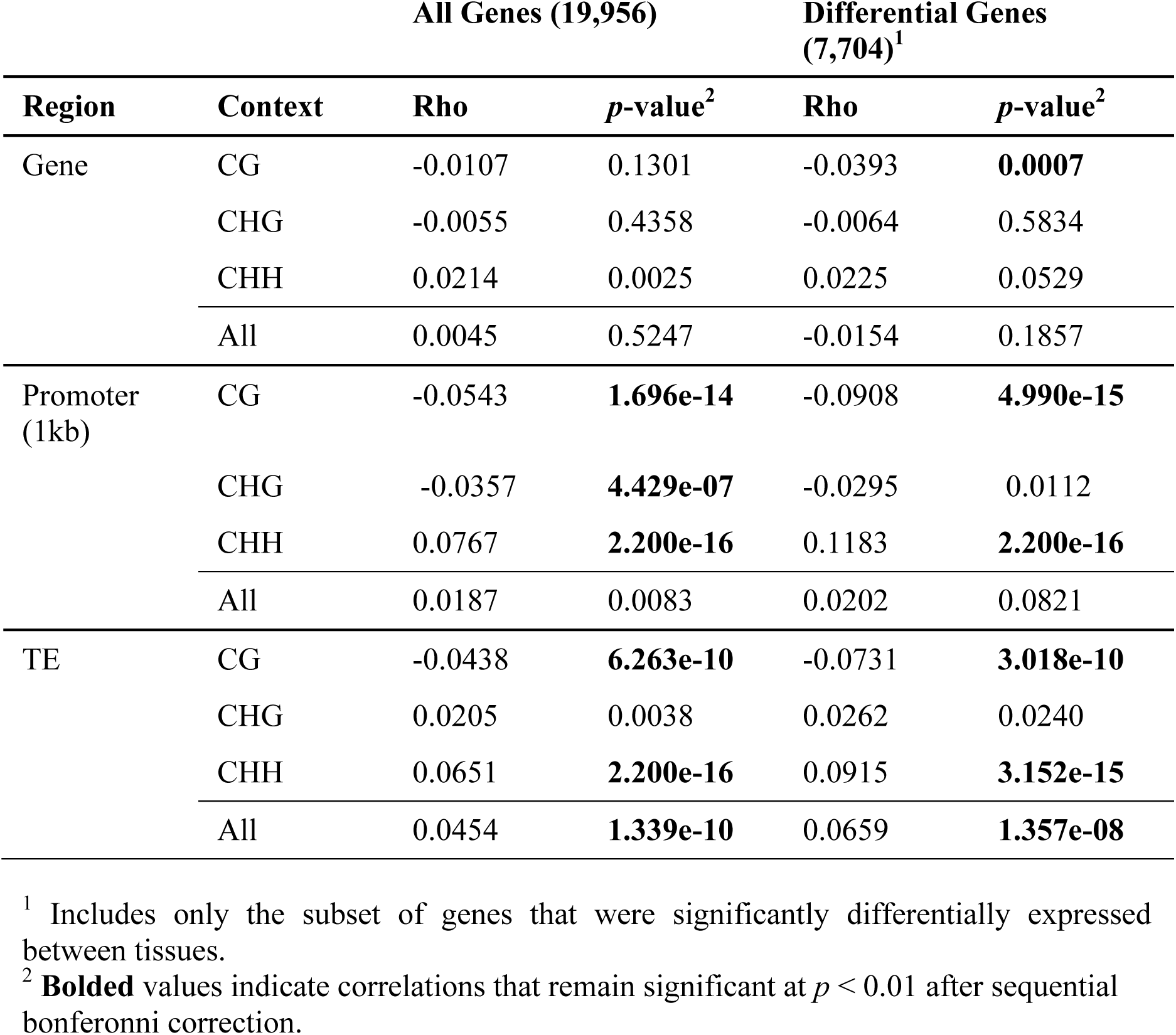
Spearman correlations between *prop* values and log2 fold change in gene expression.

#### Promoter methylation and GE

Differential methylation of promoter regions has been reported to correlate with GE during tomato ripening [42] and perhaps to tissue-specific GE of soybean genes [27]. Accordingly we assessed relationships between promoter methylation and GE. For promoter regions there is a clear expectation of an inverse relationship between methylation levels and GE [46], such that higher expression correlates with lower levels of methylation.

We first assessed the pattern of DNA methylation within promoters and note that it varies as a function of both distance from the TSS and cytosine context. For example, CG and CHG methylation both reach a zenith ∼750 bp from the TSS (Fig. 5AB), as documented previously [4,5], but CHH methylation was maximal ∼500 bp from the TSS (Fig. 5C). Within a tissue, promoters again exhibited the striking pattern of *prop*_CG_ and *prop*_CHG_ correlation, where the former is higher than the latter for 80% of observations (Fig. 5D). The same relationships was evident between CG and CHH methylation (Fig. 5F; Table 2) but not between CHG and CHH methylation (Fig. 5E).

**Fig. 5:**
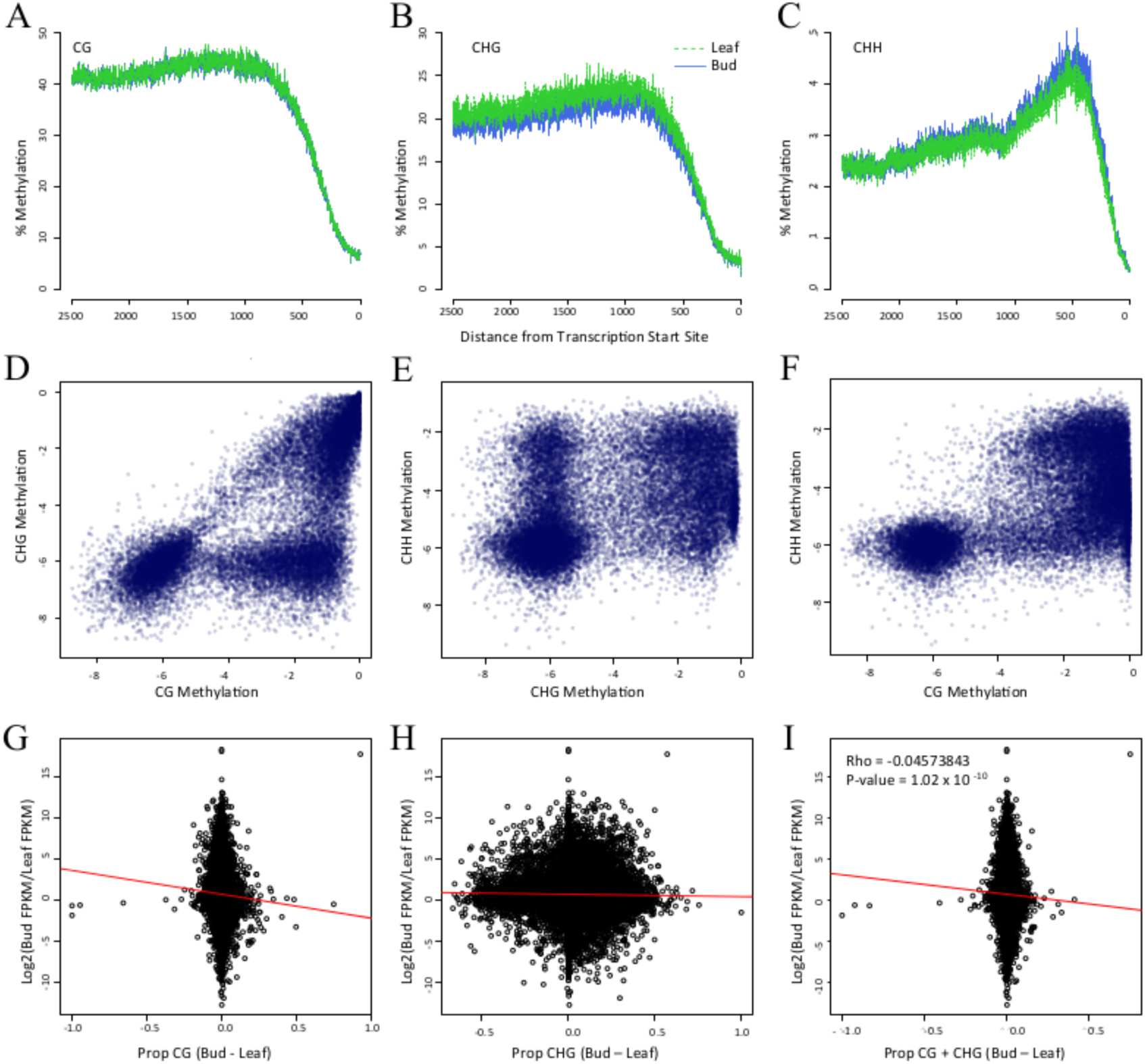
Methylation patterns within promoters and its relationship to gene expression. Graphs A,B and C present the level of CG, CHG and CHH methylation, respectively, in terms of distance from the Transcription Start Site. Graphs D, E and F compare methylation contexts, as measured by *prop* statistics in a log scale, within leaf tissue. Floral bud comparisons are not shown but are visually identical. Panels G, H and I compare differential gene expression [log2fold (FKPM_Flower/FKPM_Leaf)] vs. the difference in *prop* between floral bud and leaf tissue. The correlation values for G and H are in Table 3.

We expected a negative correlation between differential methylation and differential GE, and indeed the expected relationship was evident for both CG and CHG methylation (Table 3). With 1000 bp promoter regions, the correlation was as high as *r* = -0.0908 (p = 4.99e10^−14^) for the subset of 7,704 differentially expressed genes (Fig. 5G; Table 3). In contrast to CG and CHG methylation, *prop*_CHH_ was significantly *positively* correlated with differential GE (Table 3), showing that higher CHH methylation relates to enhanced gene expression. Overall, for promoter regions we conclude that: *i*) CG and CHG methylation covary with differential GE in the expected direction, *ii*) that CG methylation explains up to ∼9% of the variation in gene expression between tissues for differentially expressed genes, but *iii*) CHH methylation differs from the expected pattern.

#### TE methylation and GE

Because the methylation of TEs is known to suppress the expression of nearby genes [3,45,49], we expected that differences in GE would correlate negatively with differential methylation of nearby TEs. That is, if a nearest TE to a gene is more highly methylated in floral bud, we predicted it should suppress GE in flowers, thus yielding a negative correlation in our analyses. We detected this predicted negative correlation but only in the CG context (Table 3). In contrast, correlations between differential GE and both *prop*_CHG_ and *prop*_CHH_ were positive, with the *prop*_CHH_ comparisons reaching statistical significance (Table 3). Across all contexts (*prop*_C_), the relationship was also significantly positive, likely owing to the positive trends for *prop*_CHG_ and *prop*_CHH_ countervailing the trend for *prop*_CG_. Finally, we also applied a linear model to disentangle the effects of methylation vs. the distance (in bases) of the TE from the gene (S4 Table). In the linear model, the effect of methylation remained significant (*p* < 10^−3^), but the effect of distance explained little and was not significant.

## Conclusions

Although there is a widespread belief that methylation affects gene expression during development [1], relatively few studies have contrasted methylation and gene expression between tissues on a genomic scale. Moreover, BSseq data have rarely been replicated in these studies. Hence, our first goal was simple: to determine whether methylation between two tissues is, in fact, differentiated beyond the level expected from proper replication. For this comparison we chose two tissues that have been sampled commonly in other plant studies - leaves and floral buds. Overall, we were able to detect a significant signal of differentiation between tissue samples based on two methodological approaches (permutation tests and clustering analyses) and two measures of variation (DMSs and DMRs).

Nonetheless, a sobering observation was that the false positive rate (FPR) was extremely high for contrasts between single replicates. For DMS analyses, the lowest FPR in our analyses was 75%. In other words, had we based our inferences on single replicates, three-quarters of our inferences about the sites of “tissue-specific” methylation would have been incorrect relative to inferences based on the larger, replicated dataset. The FPR for DMR analyses was similarly large, at least 80%. While there are ways to decrease the FPR statistically in theory, they may result in the cost of sensitivity and power. Such tradeoffs in the use of BSseq replicates are the topics of ongoing theoretical and algorithmic research, but thus far these render improvements only for data with less coverage those in this study [50]. Altogether, we conclude that reliance on single BSseq replicates may be misleading when the goal is to focus on specific DMRs or DMSs. For this reason, we recommend analyzes that summarize over a region – e.g., genes [48] or promoters or TEs – as opposed to individual sites or individual DMRs. Moreover, because replication has been applied so rarely in plant studies, we hope that our description of within-and between-tissue replicates helps guide interpretation of the existing literature.

Although we detected significant methylation differentiation between tissues, our results were similar to previous studies in documenting that tissues are far more similar than different in their methylation patterns [22]. For example, we detected ∼37-fold more sites conserved between tissue samples than variable sites. Most of the observed differences occurred in the CHG context within TEs and promoters, but there were also slight biases in *total* methylation between leaf and floral bud. Overall, these observations add to the growing notion that methylation differences between plant tissues are slight, except for a few exceptional tissues, such as the endosperm and the pollen vegetative nucleus [19-22]. Since neither of these tissues contributes to ensuing generations, these epigenetic changes may be of little evolutionary consequence, although it seems that the pollen vegetative nucleus may play a role in generation-to-generation epigenetic reprogramming [23].

We have shown that tissue-specific methylation differentiation is higher than variation among replicates, but do any of these methylation differences drive functional differentiation? To address this question, we generated RNAseq data for the same sets of replicates and examined the correlation between differential GE and differential methylation, many of which were significant. The most striking aspect of these results is that they vary by methylation context. In general, CG methylation correlates with GE as predicted: higher CG methylation in one tissue correlates with lower GE in that tissue. This relationship is true whether one examines genes, promoters or TEs (Table 3). In contrast, the results for CHH and CHG variation are more varied, with CHH methylation trending in the opposite direction than predicted for both promoters and TEs. These observations indicate that CG methylation is the primary component of variation to affect (or at least covary with) GE. This observation is consistent with the fact that genic expression in pines covaries with CG but not CHG methylation, even though pine genes are heavily methylated in both contexts [47].

Another interesting aspect about methylation contexts is that they appear to be hierarchical, because typically neither CHH nor CHG variation exceeds CG methylation, regardless of the region under consideration (Figs. 4A, 5DEF). These results suggest that CG methylation acts in some unknown way to limit methylation in the other contexts, at least in brachypodium. It remains to be seen whether this relationship holds for other species and additional tissues.

Overall, our study suggests that methylation patterns covary with tissue-specific expression, but also that differential CG methylation explains only a small proportion of tissue-specific variation in GE (i.e., between 1% and 9% of variation; Table 3). We note, however, that our study likely underestimates the magnitude of the effect, for at least two reasons. First, the predictive power will probably increase with the number of tissues sampled. An explicit goal of future studies should be to estimate the percentage of GE variation explained by DNA methylation based on a broader range of tissue-types; however, to do so will require better sampling – both in terms of tissues and replicates – than has been performed to date. Second, like most other papers in plant epigenetic research, our tissue samples undoubtedly included multiple cell types; indeed we suspect that the variation in cell types is the primary reason for high variation in DMSs and DMRs among biological replicates (Fig. 1). A recent review has called to question the value of ‘tissue’ vs. ‘cell’ samples [40]. In the review, the authors argue that the signal of differentiation for highly specialized cells will be masked within tissue samples that contain multiple cell types. This may or not be true, as it depends critically on the as-yet-unknown pattern of cell differentiation and of course the cellular composition of tissue samples. Nonetheless, their point is well taken: it is possible that tissue, as opposed to cell-type, samples lead to underestimate of the overall contribution of epigenetic variation to gene expression.

## Materials and Methods

### BSseq Data and Mapping

The BSseq data were published previously [35] and were available in the Short Read Archive (accession nos. SRX208151–SRX208156). Briefly, three *B. distachyon* plants from the Bd21 line were grown under 20-h days to induce rapid flowering. Spikes and leaves were harvested at the beginning of anthesis. For each plant and tissue, ∼two micrograms of genomic DNA was sonicated and purified using Qiagen DNeasy mini-elute columns (Qiagen). Sequencing libraries were constructed with the NEBNext DNA Sample Prep Reagent Set 1 (New England Biolabs, Ipswich, MA) but with methylated adapters in place of the genomic DNA adapters. Ligation products were purified with AMPure XP beads (Beckman, Brea, CA). DNA was bisulfite treated using the MethylCode Kit (Invitrogen, Carlsbad, CA) following the manufacturer’s guidelines and then PCR amplified using Pfu Cx Turbo (Agilent, Santa Clara). Libraries were sequenced using the Illumina HiSeq 2000. The BSseq reads were mapped to the brachypodium reference genome (version 1.0) following [35], which included filtering of low-quality reads and bases (q < 20) and mapping with BRAT software [51]. Mismatches for mapping were allowed only at potentially methylated sites.

### mRNAseq data and analysis

RNAseq data were generated from the same tissue samples as BSseq [35]. RNAseq relied on total RNA isolation with the Qiagen RNeasy Kit, cDNA generation with the Ovation RNA-seq system v. 2 and library preparation with the Illumina TruSeq DNA Sample Prep. V2. The libraries were sequenced on the HiSeq2500 (100 cycle, single read) in the UCI High Throughput Genomics Facility. RNAseq reads were processed using Trimmomatic (v 0.30) to remove low quality reads (<20) and adapter sequence.

Analyses of RNAseq data was based on read mapping to the *B. distachyon* MIPs v.1.2 reference sequence, using TopHat (v1.49.0) [52] with default parameters. In this analysis, reads were counted for each annotated gene, so long as that gene did not overlap with an annotated transposable element (see below). Reads were counted for each gene in each replicate, and then DESeq (v1.16.0) [53] was employed to identify differential expression between tissues with a false discovery rate of *q* < 0.01. For the comparison of differential gene expression and differential methylation we used all genes that had the number of fragments per kilobase of transcript per million mapped reads (FPKM) > 0.02 in both tissues. The difference in gene expression was defined as [log2 (Flower_FPKM)/(Leaf_FPKM)], where Flower_FPKM and Leaf_FPKM were based on data from all three replicates.

### Definitions of genomic features, DMSs, CMSs, and *prop* values

We used genome annotations to define genes, promoters and TEs. A gene was defined from the transcription start site (TSS) to the transcription stop site, including putative introns, using the MIPs (v1.2) annotation [54]. TEs were also based on the MIPs (v1.2) annotation. TEs that overlapped with genes were removed from analysis along with any genes that were contained in a TE. Gene annotations were also the basis for promoter annotation, which were defined as the 1.0 kb region upstream from the TSS.

To determine whether individual cytosines were methylated or unmethylated, we computed a binomial probability at a significance level of *p* ≤ 0.01, following [5]. This probability required a rate of conversion error, which was calculated on contaminating chloroplast data and was ∼1% [35]. The specific error rate for each tissue was found for each replicate and for each tissue (i.e., L1+L2+L3 and F1+F2+F3; S1 Table). Once a base was defined as methylated or unmethylated in each tissue, a base that was methylated in each tissue was deemed a conserved methylation site (CMS).

To identify Differentially Methylated Sites (DMSs), we applied a Fisher exact test (FET), which was based on a 2X2 table of the number of converted Cs to non-converted C’s across the two samples [5]. A site was considered as differentially methylated between two samples – i.e., a DMS – when the FET yielded a p-value < 0.05.

DMRs were defined by the number of DMSs in a row that had a consistent direction of methylation bias (i.e., hypermethylation in leaf or flower), that were within 500 bp of each other and that were uninterrupted by a CMS or by a DMS in the opposite direction. We considered DMSs in all contexts (i.e., CG, CHG and CHH) to define DMRs. To assess significance, we calculated the length of DMRs (as defined by the number of unidirectional DMSs) expected to be found at random in the genome, given both the underlying distribution of cytosines in proper context and the numbers of DMSs and CMSs. To calculate the random expectation, we permuted DMSs and CMSs among genomic sites in their appropriate contexts, identified DMRs within permutated genomes, and ascertained DMR lengths. After permuting across the genone, we identified DMRs and noted the number of DMSs that constitute each DMR (S1 Fig.). DMRs that were of a length expected found at *p* < 0.01 in the permuted genome were considered ‘significant’ for analysis of observed data.

The final metric was the proportion of methylation or *prop*_*c*_, which was used as a measure of methylation across a region. The *prop* value was determined by adding the total number of converted reads over the total number of reads for cytosines in a specific context. The context could be CG (*prop*_CG_), CHG (*prop*_CHG_), CHH (*prop*_CHH_) or all three contexts (*prop*_C_).

### Additional Statistical Analyses

To construct the trees in Figure 1, distance values were converted to Newick format and unrooted neighboring-joining trees were made using the ape and phyclust libraries in R [55].

To determine whether DMRs were closer to differential expressed genes than other genes, we first labeled each gene as either differentially expressed or not. We then calculated both the observed distance from a differentially expressed gene to its closest DMR and its average across the genome. We then randomized the labels (differentially expressed or not) among genes within the genome and recalculated the average distance between a differentially expressed gene and its nearest DMR. The randomization was performed 1000 times to generate a distribution of the average distance from a DMR to a gene and to determine whether the observed average was extreme (S4 Fig.).

All correlations were based on cor.test in R, using the Spearman correlation.

## Acknowledgements

We are grateful to D. Garvin (USDA, Minnesota) for plant material and to R.L. Gaut for generating the BSseq and RNAseq dataset. K. Roessler was supported by the National Institute of Biomedical Imaging and Bioengineering, National Research Service Award EB009418 from the University of California, Irvine, Center for Complex Biological Systems. S. Takuno is supported by an internal grant from SOKENDAI and JSPS Grant-in-Aid for Young Scientists (B) (Grant No. 15K18585). The work was supported by NSF grant IOS-1542703 to BSG.

## Supporting Information

**S1 Fig. A histogram of the length of DMRs found after randomization of methylated cytosines within the brachypodium genome.** Methylated cytosines were randomized in the proper context, and the number of DMRs in the same direction were counted. Within a randomized genome, a run of five or more methylated cytosines in length represented 1.3% of all potential runs; we defined a DMR to be ≥ 5 methylated cytosines in the same direction, because this length represented a significant observation at the p ∼0.01 threshold. See Materials and Methods for additional details.

**S2 Fig. Plots of chromosomal densities of methylation features.** Plots of chromosomal densities of A) CMSs, B) DMSs, C) genes, and D) TEs. Density was measured within a 50kb sliding window for smoothing. E) The graphs plot differential gene expression plotted along the physical length of chromosomes. This figure mimics Figure 2 of the main text, but includes the remaining four chromosomes.

**S3 Fig. A volcano plot of the 26,072 genes tested for differential gene expression between leaf and floral tissue samples.**

**S4 Fig. A histogram of the average distance between DMRs and genes**. The histogram is based on 1000 randomizations (see Materials and Methods). The red line denotes the observed value.

**S1 Table: The estimate of the rate of conversion error**. Estimates are based on analysis of chloroplast DNA in each replicate or on combined replicates.

**S2 Table: A summary of RNAseq data.** The table provides the number of reads after quality trimming, the number of reads that TopHat used to map for both left and right reads, and the maximum and minimum read lengths. The total number of transcripts is from based on output from cufflinks.

**S3 Table: Go enrichment terms for differentially expressed genes between leaf and flower**. Only the enrichments terms with a p-value < 0.01 are given.

**S4 Table: Results of the application of linear models**. The models test for an effect of the TE distance to a gene and of TE methylation to differential gene expression.

